# The Impact of TLR9 Expression Loss on Breast Cancer Tumor Cells

**DOI:** 10.1101/2024.07.04.601915

**Authors:** Emily Sible, Gregory Weitsman, Salome Amoyal, Guillaume Roblot, Marie Marotel, Rémi Pescarmona, Nathalie Bendriss-Vermare, Cheryl Gillet, Amie Ceesay, Marie Cecile Michallet, Christophe Caux, Francois-Loic Cosset, Umaima Al Alem, Tony Ng, Uzma Ayesha Hasan

**Affiliations:** Centre International de recherche en Infectiologie, CIRI, Inserm, U1111, Lyon, France; King’s College London, London, UK; Research Oncology, Division of Cancer Studies, Guy’s Hospital, King’s College London, London, UK; Cancer Research Centre of Lyon, CRCL, INSERM U1052-CNRS UMR5286, Lyon, France; Division of Epidemiology and Biostatistics, University of Illinois at Chicago, Chicago, Illinois, USA

## Abstract

Toll-like receptor 9 (TLR9), primarily expressed in human dendritic and B cells, recognizes double-stranded DNA motifs from pathogens, initiating an inflammatory response. Recent studies have revealed TLR9’s involvement beyond its conventional role in immune response, notably during the tumorigenesis of various cancers such as head and neck, cervical, and ovarian cancers. Here we show by immunohistochemistry analysis demonstrated significantly lower TLR9 levels in breast cancer tumors compared to normal breast tissue epithelium. Similarly, TLR9 downregulation was also observed in several transformed breast cancer cell lines versus untransformed breast epithelial cell lines. Furthermore, overexpression of TLR9 led to reduced proliferation potential of breast cancer cell associated with activation of senescence as was evident by upregulation of proinflammatory cytokine IL-6, chemokines IL-8 and CXCL1; and growth factor GM-CSF. These findings support TLR9’s regulatory role in mitigating breast cancer and highlight its critical connection between the inflammatory response and tumor immunity.

## Introduction

Breast cancer (BC) is the most frequent malignant disease in females worldwide [1], where 1 in every 6 cancer case resulted in death as of 2020 [2]. Current models predict an ever increasing global burden, particularly amongst premenopausal and postmenopausal women susceptible to age-related risk factors [3]. Cancer of the breast tissue occurs when transformed epithelial cells-if not removed-invade the breast duct (also known as invasive ductal carcinoma), given the proximity to the lymphatic system and blood stream the risk of metastasis to other tissues is high and treatment efficacy is relatively low [4]. The incidence of invasive ductal carcinoma can be dramatically reduced by improving our abilities to detect, diagnose, and treat preinvasive patients. However, the transformed epithelium exhibits intratumoral heterogeneity and molecular diversity not only within the same individual [5,6], but in presentation between patients [7,8], which presents an informational challenge for clinicians and researchers alike. Indeed, immune cells have been shown to infiltrate early BC lesions [9]. Yet the exact mechanisms through which the body’s innate immune system contributes to the process of carcinogenesis are not fully understood.

The Toll-like receptor (TLR) family are pattern recognition receptors that respond to pathogen-associated molecular patterns and damage-associated molecular patterns, canonically important in the innate immune response [10]. TLR9 is a sensor specialised in recognizing double stranded DNA (dsDNA) motifs expressed by pathogens or released from damaged host cells. In humans, TLR9 is expressed mainly on immune cells such as macrophages, plasmacytoid dendritic cells, and B or T cells [11], but can also be found in the non-immune compartment such as epithelial cells in the mammary gland [12,13] or even breast milk [14]. Activation of TLR9 which leads to increased production and secretion of inflammatory cytokines and chemokines, particularly type I interferon, a critical mediator involved in tumor immunosurveillance and rejection [15,16]. Importantly, changes in the secretory phenotype of these inflammatory factors (i.e. IL-8, IL-6, CXCL-1) are associated with cellular senescence and growth arrest of malignant cells [17,18], indicating an important bridge between inflammation and tumor suppression. In addition to immune activation, TLRs have previously described roles in tissue repair during epithelial homeostasis or after injury [19,20]. Additionally, *in vitro* data from cervical and head and neck cancer demonstrated that overexpression of TLR9 induced a delay in S-phase exit and upregulation of tumor suppressor proteins (i.e. p16, p21, p27, and p53) [21].Collectively, these data suggest TLR9 may play a central role in modulating the tumor microenvironment during carcinogenesis. Furthermore, TLR9 has been shown to be under-expressed in the epithelium of many cancers [22–25], which may introduce important context for the role of TLR9 in the transformed epithelium during breast cancer metastasis.

In breast cancer, the role of TLR9 is not fully understood, and research findings are conflicting. Some studies suggest that TLR9 expression is increased in breast cancer cells, and its activation may contribute to the promotion of tumor growth and aggressiveness [25–28]. On the other hand, other studies propose that TLR9 overexpression can have anti-tumor effects by stimulating immune responses against cancer cells [29–31]. Here, we observed a downregulation in TLR9 expression in clinical BC samples and human breast cancer cell lines. Furthermore, TLR9 overexpression in BC cells induces a reduction in cell growth concurrent with an increase in markers of cellular senescence. Thus, TLR9 expression may control the cell cycle to repress cell growth and detain the transformation of epithelial cells during BC development.

## Results

### TLR9 is differentially expressed in breast cancer epithelium

To investigate if TLR9 protein can be specifically identified in the breast epithelial cells, we optimized the TLR9 protein staining on skin, human tonsil, and healthy breast tissue. We observed weak membranous staining in skin tissue in the keratinocytes and tonsillar lymphocytes. The luminal cells exhibited diffuse cytoplasmic staining patterns in normal breast tissue with minimal background signal interferences (**Figure 1)**. To further test the specificity of the primary antibody, we utilized HEK293T cells that were stably expressing TLR9. TLR9 staining was effectively inhibited upon the use of blocking recombinant TLR9 peptide (**Figure 2**).

**Figure 1:**
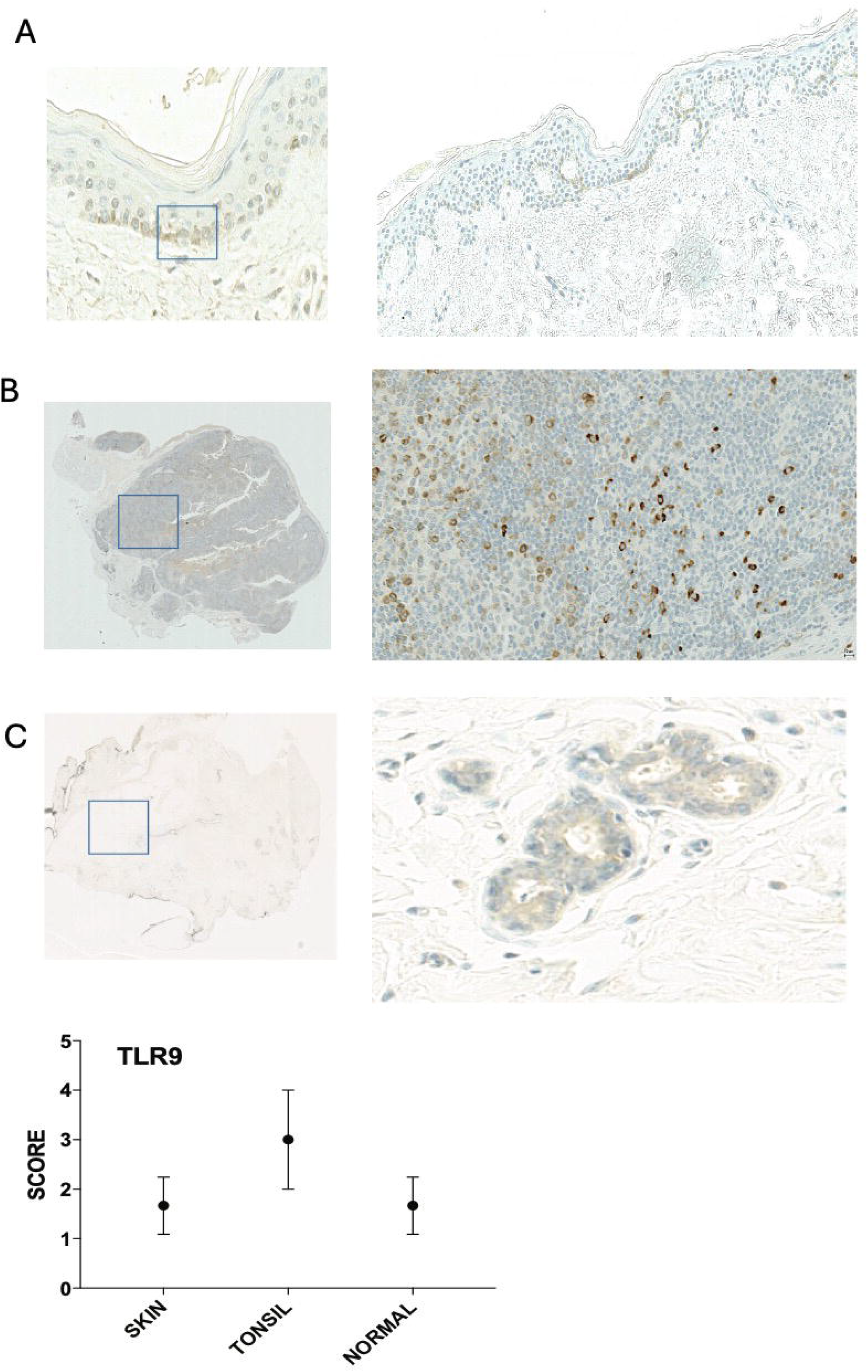
IHC Scoring of TLR9 in epithelial tissue immunohistochemical analysis of TLR9 expression on paraffin-embedded skin (A), tonsil (B), and breast tissue (C).

**Figure 2:**
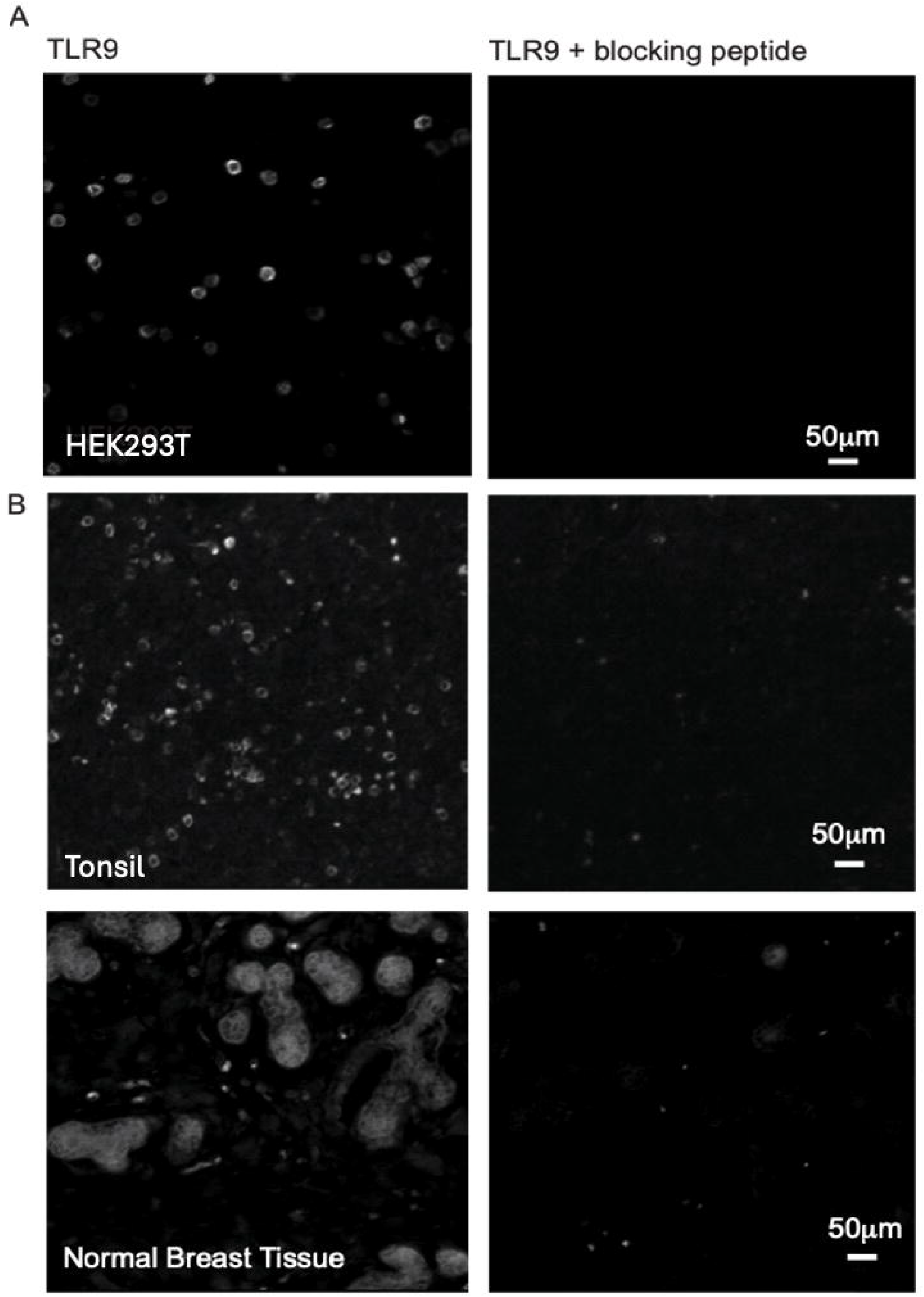
Specific staining of TLR9 observed in breast epithelial tissue immunofluorescent staining of HEK293T (A), tonsil (B), and breast tissue (C) samples with primary TLR9 antibody, which is effectively blocked upon use of TLR9 inhibitory peptide (right).

To determine how TLR9 is expressed in the breast epithelium, we examined both whole tissue sections and TMA (tissue microarray) spots from paraffin-embedded tumors and normal breast tissue samples. After optimizing immunohistochemistry (IHC), we stained formalin-fixed and paraffin-embedded breast tissues obtained from King’s College Tissue Bank, UK. We randomly selected a sample size of 98 breast tissue specimens, consisting of 29 cases of normal breast tissue, 39 cases of peritumoral breast cancer tissue, and 29 cases of invasive breast cancer tissue along with their adjacent tissues **(Figure 3)**. Adjacent tissue refers to the sections of tissue surrounding the breast cancer tissue in the whole tissue sections, whereas peritumoral tissue refers to the noncancerous epithelial tissue adjacent to the tumor in the same TMA spot. TLR9 expression to contextualize its spatial and temporal expression in patient-matched samples were scored.The results were evaluated by one-way ANOVA. We observed a statistically significant difference among the breast tissue samples (normal, adjacent, peritumoral and tumor tissue, p<0.0001). Tukey’s Test for multiple comparisons found that the mean value of TLR9 was significantly different between normal tissue and all other breast tissues. Additionally, the staining percentage in tumoral samples and the adjacent epithelium was on average half that observed in healthy tissues (**Figure 3A)**, suggesting an overall trend towards decreased expression of TLR9 in BC tissue, consistent with the phenotype observed in cervical cancer [32]. Furthermore, the mean TLR9 immune response score was significantly lower in tumoral specimen compared to normal epithelium (**Figure 3C**, p<0.007). Strikingly, 100% of the cells scored from TMA samples were negative for TLR9. Collectively, these data suggest that TLR9 expression is suppressed in BC.

**Figure 3:**
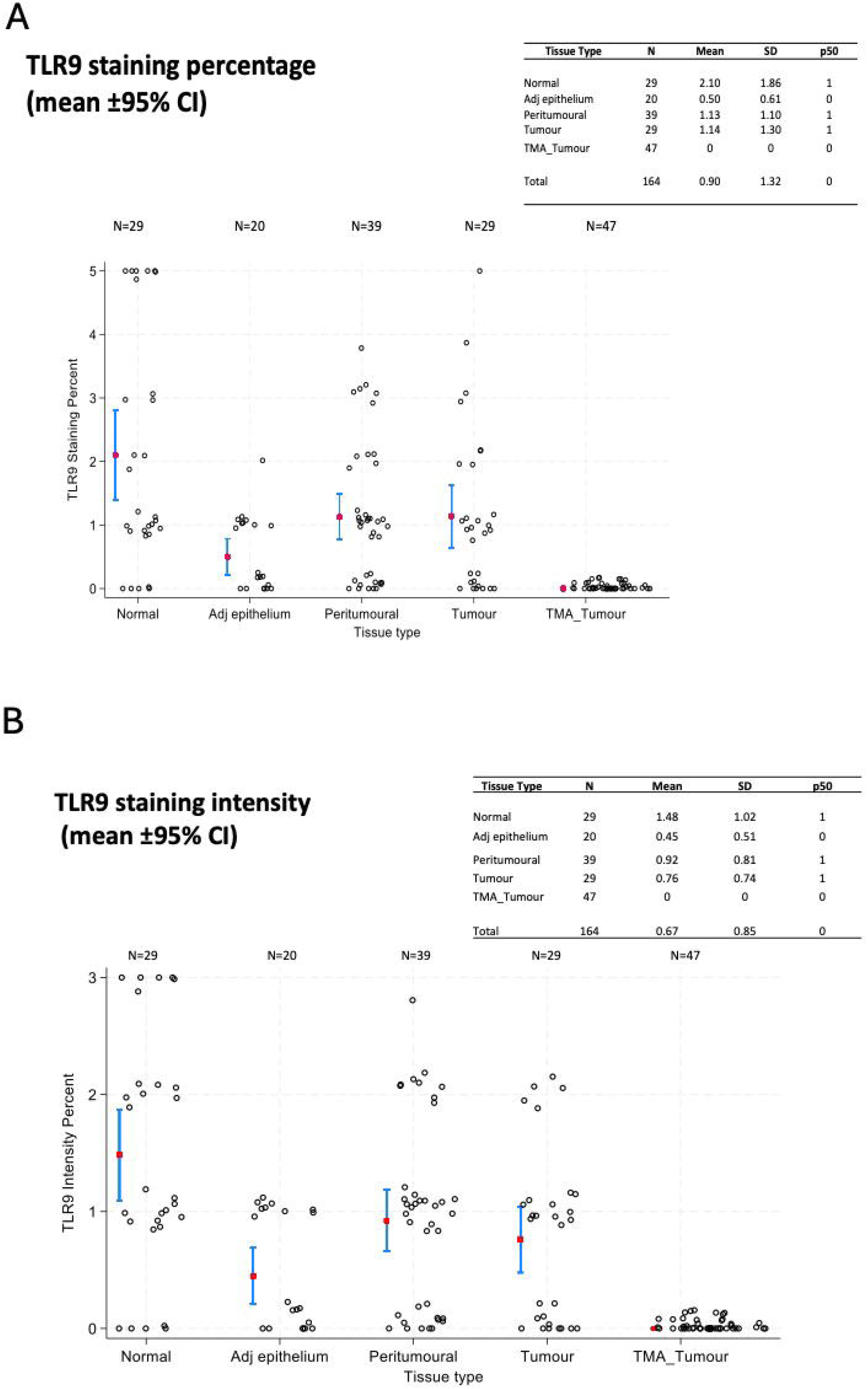
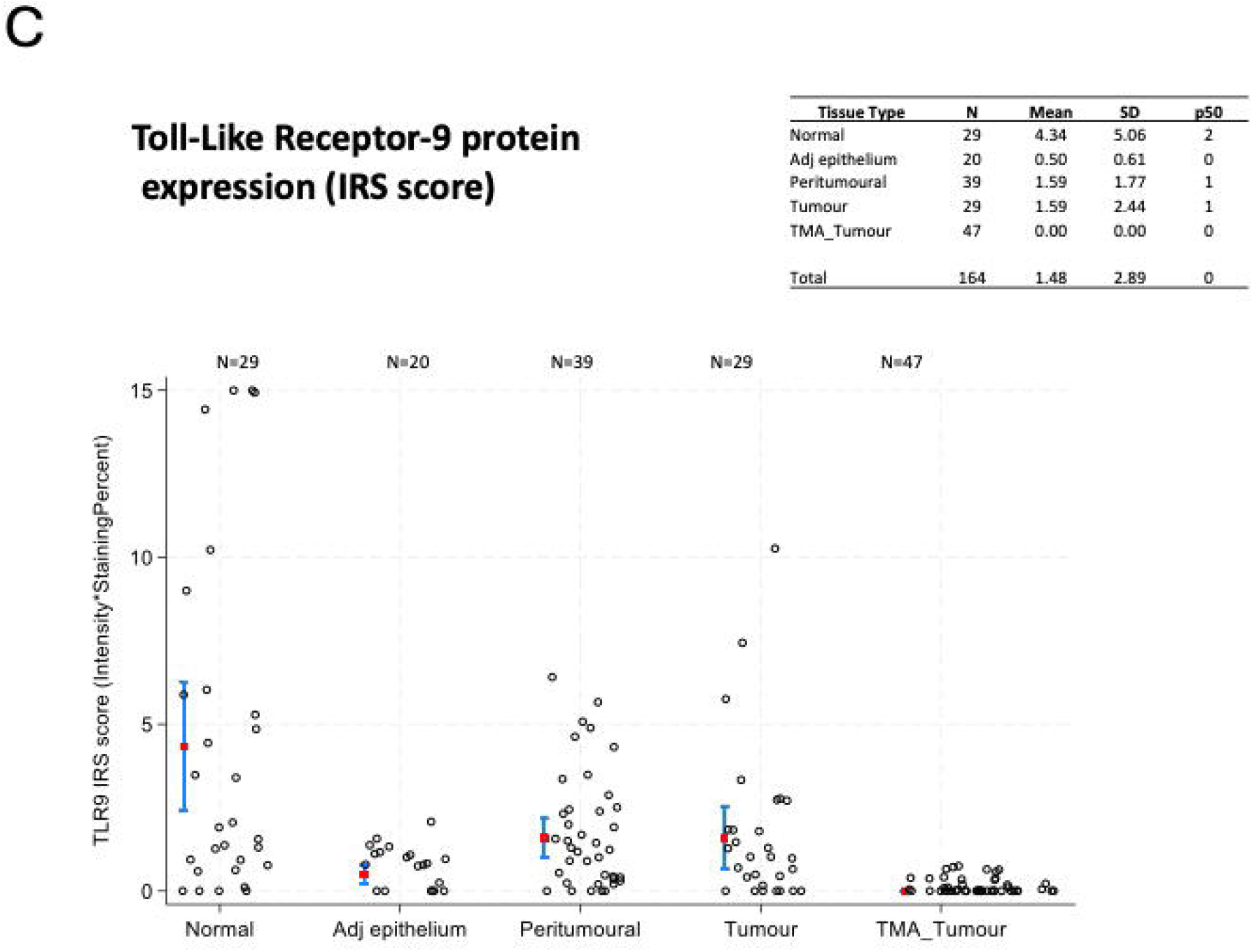
Reduced TLR9 expression in tumor microarray (TMA) samples from breast cancer patients TLR9 expression in human breast cancer samples. Statistical analysis by one way ANOVA. All scores were graded in 29 Normal breast tissue, 39 peritumoral, 24 adjacent normal epithelial, 29 tumors, and 49 TMA tumor breast tissue samples. (A.) Mean TLR9 staining percentage (±95% CI). The mean TLR9 expression was significantly lower in the tumor specimens compared with normal epithelial cells (p < 0.0001). (B.) Mean TLR9 staining intensity (±95% CI). The mean TLR9 expression was significantly lower in the tumor specimens compared with normal epithelial cells (p < 0.0001). (C.) Mean TLR9 immune response score (positivity*intensity) (±95% CI). The mean TLR9 expression was significantly lower in the tumor specimens compared with normal epithelial cells (p < 0.007).

### TLR9 expression is reduced in transformed breast cancer cell lines

To investigate if the phenotype in primary tumors is observed in transformed cells, we mined microarray expression data in 59 BC cell lines from the Cancer Cell Line Encyclopedia (CCLE). Indeed, we observed a significantly lower expression of TLR9 (p <0.0001, Mann-Whitney) in all BC cell lines compared to immortalized cell lines (n=8) (**Figure 4A**). Importantly, the immortalized breast cells HMEL showed the highest expression of TLR9 (**Figure 4A**). We validated the *in silico* analysis in six cell lines (MDA-MB-231, SK-BR-3, Hs578T, HCC1937, T-47D, and MDA-MB-361) by quantitative PCR and found that relative TLR9 mRNA levels were lowest in the MDA-MB-361 cells (**Figure 4B, 4C**). Collectively, these data support the clinical observations and support the conclusion that TLR9 expression is lost during breast carcinogenesis.

**Figure 4:**
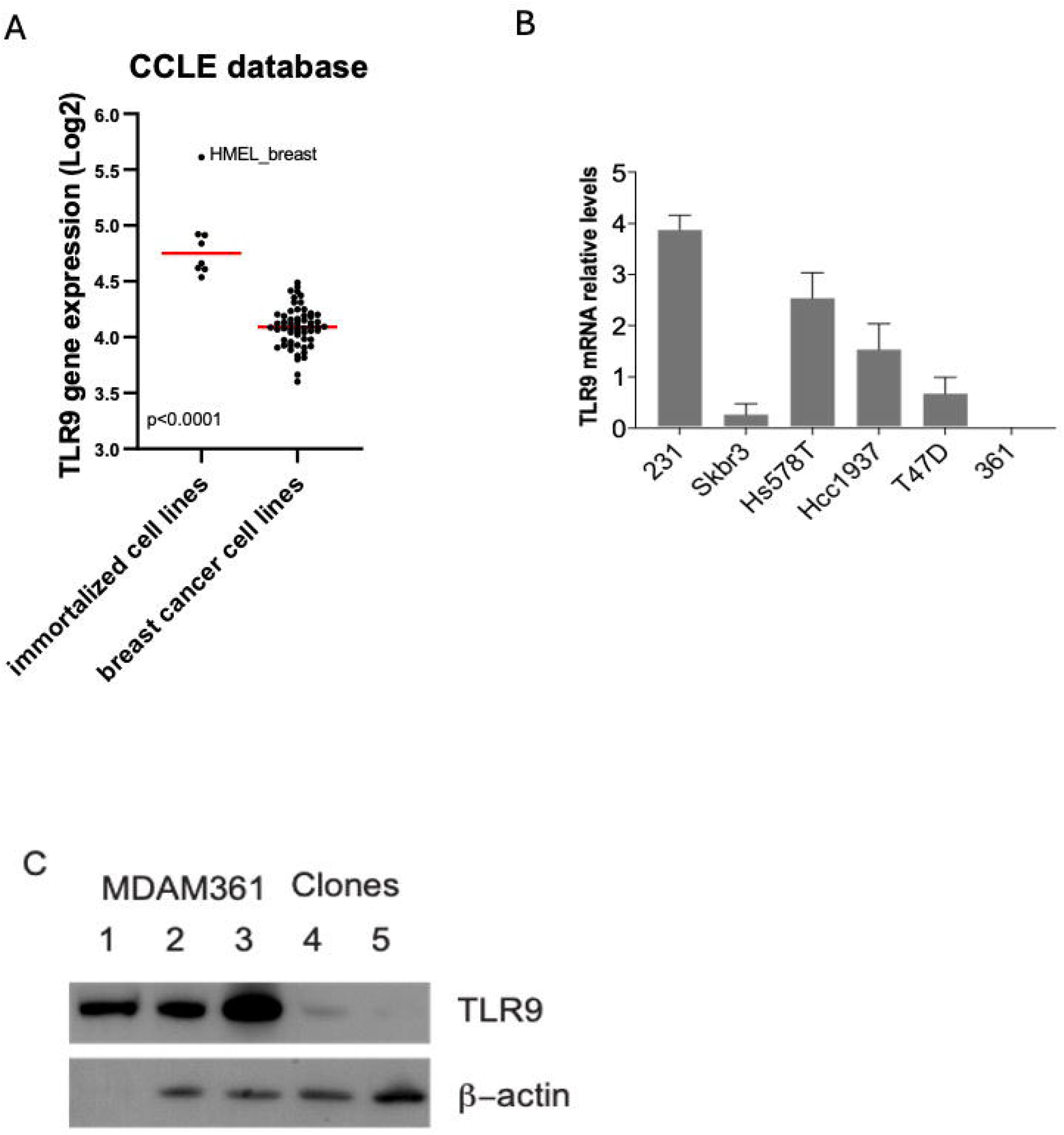
TLR9 in BC cell lines. (A) TLR9 gene expression analysis was generated by using Array Studio software (Omicsoft Corporation). CEL files of BC (N=59) and immortalized (N=8) cell lines were downloaded from the Cancer Cell Line. Each symbol represents an independent breast cancer cell line or immortalized cell line. (B) Quantitative PCR of cDNA from six BC cell lines. (C) Western blot analysis of TLR9 clones generated using the MDAM361 cell line.

### TLR9 impairs cell proliferation and induces senescence in MDAM-MB-361 BC cells

To examine the mechanisms underlying TLR9 downregulation during breast carcinogenesis, we transduced MDA-MB-361 cells to overexpress TLR9 with a doxycycline-inducible *Tet-*on lentiviral vector pLVUT carrying a transgene for TLR9 or TLR7 as a control (**Supplemental Figure 1**). Upon treatment with doxycycline, TLR9 competent MDA-MB361 grew significantly slower in culture over time than TLR9-cells (**Figure 5A**, p<0.005). To assess the clonogenic potential in BC cells, we examined colony formation of TLR9+ and TLR7+ MDA-MB361 cells upon induction with 10uM, 100uM, and 1000uM doxycycline (**Figure 5B**). Notably, we observed a significant reduction in colony number (**Figure 5C**, p<0.005) as well as size (**Figure 5D**, p<0.005) in TLR9+ cells compared to TLR9-cells, regardless of doxycycline dosage. Furthermore, no significant growth defects were observed in TLR7 controls (**Figure 5C, 5D**). To ascertain if TLR9+ MDA-MB361 cells were undergoing cell cycle arrest, cells were stained for Edu and analyzed by flow cytometry (**Figure 6**). We observed that TLR9 overexpressing cells had fewer cells (22.3%) in S phase compared to the TLR9- or TLR7 controls (approximately 35%), suggesting that fewer TLR9+ cells are progressing through the cell cycle. These data suggest that TLR9 induces a reduction in cell proliferation in BC cells.

**Figure 5:**
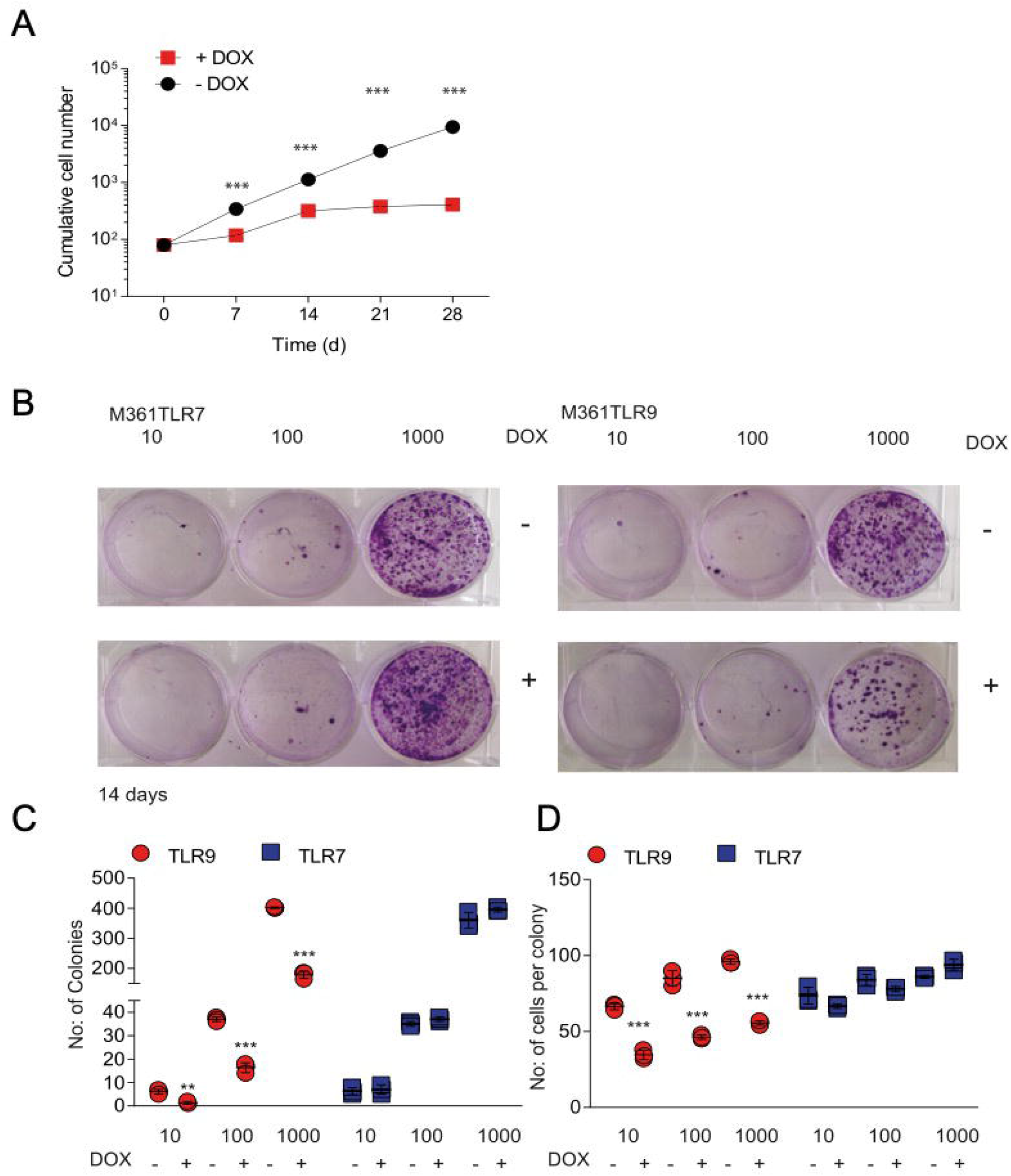
TLR9 reduces cell number and colony formation *in vitro*. (A) Proliferation curve in TLR9 proficient (+dox) and TLR9 deficient (-dox) cells. Reduction in number of colonies (B) and cells/colony (C) in TLR9+ (+dox) compared to TLR9-(dox-) cells (p <0.005) at various doxycycline concentrations. No significant growth defects occurred in TLR7 controls. Representative image of colony assay at day 14 is shown (D).

**Figure 6:**
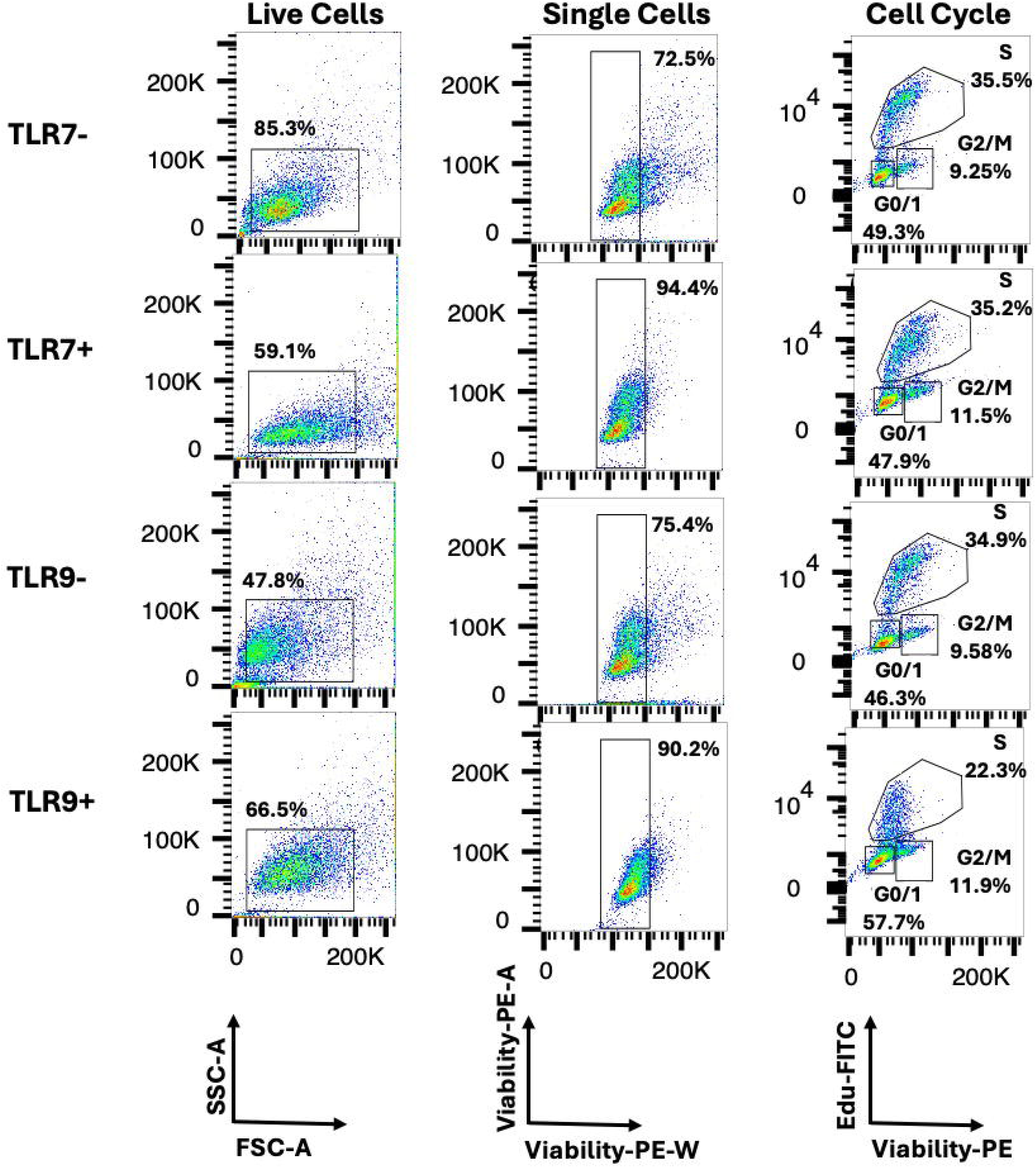
TLR9 overexpression abates progression through S phase. Flow cytometry analysis of Edu incorporation (FITC). Cells were stained with propidium iodide (PE) to assess for viability.

To understand if the arrest in cell growth was induced by cellular senescence, we stained for senescence-associated β-galactosidase (SA-β-Gal) as described previously [18]. Strikingly, TLR9-expressing MDA-MB-361 cells appear more positively stained than TLR9-cells (**Figure 7A**). Furthermore, TLR7 expression did not affect SA-β-Gal staining. A key feature of senescent cells is the production of secretion associated senescent proteins (SASPs)-comprised of various chemokines, cytokines, and growth factors-which induce a “pro-senescent” microenvironment through autocrine and paracrine signaling [17]. To examine if TLR9 expression in BC cells induces a change in SASP components we cultured pLVUT-TLR9 or pLVUT-TLR7 MDA-MB-361 cell in doxycycline for 3-4 days, then harvested the supernatant to incubate with a cytokine array as described previously [18]. Notably, the TLR9+ BC cells exhibit an increase in hallmark SASP components, particularly IL-8, CXCL1, GM-CSF) (**Figure 7B, Figure 7C**). Although the cytokine assay revealed moderate differences in the IL-6 family (IL-6 and oncostatin M), we found the relative mRNA expression of IL-6 to be higher in TLR9+ than TLR7+ cells (**Figure 7D**). Collectively, these data indicate that senescence is induced in BC cells in a TLR9 dependent manner.

**Figure 7:**
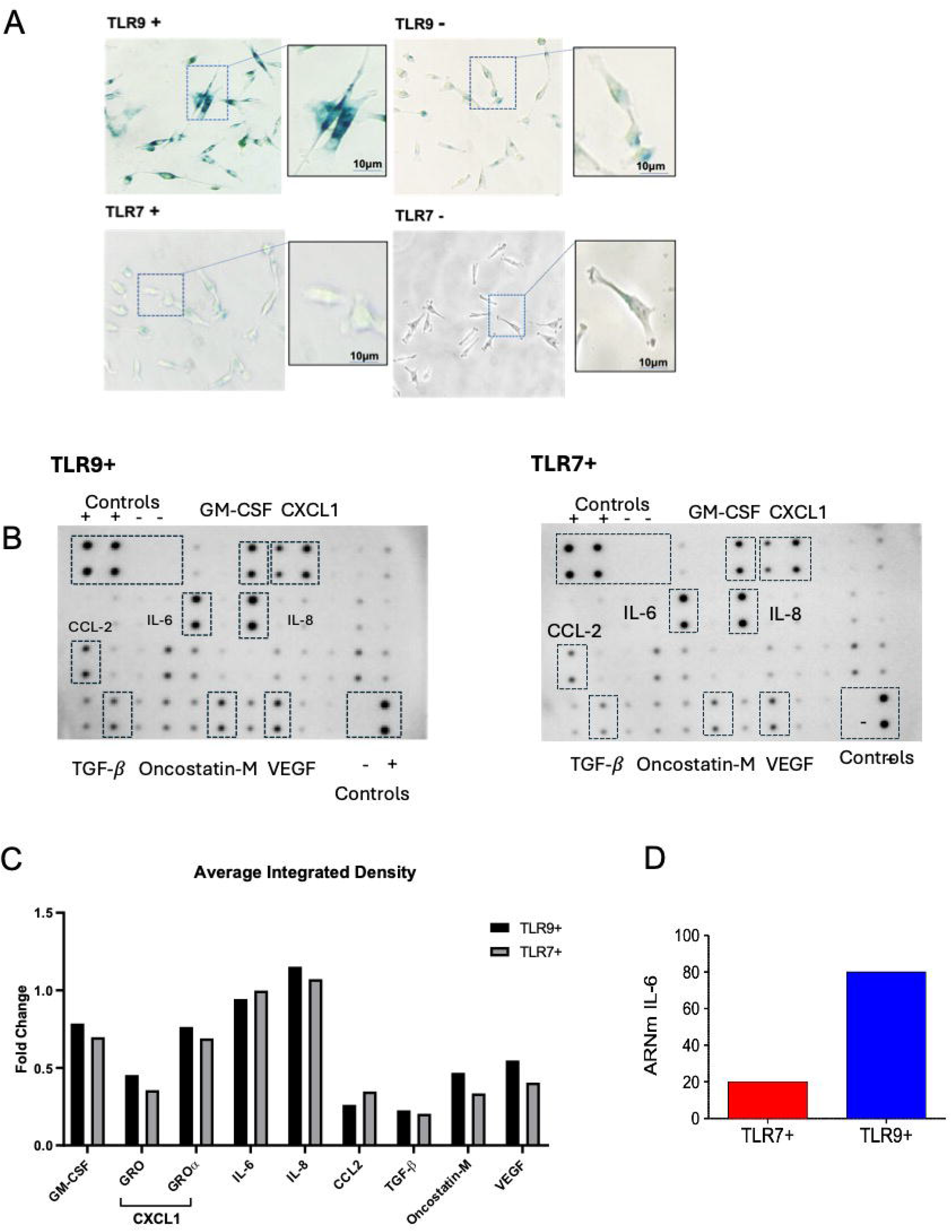
TLR9+ BC cells exhibit markers of senescence. (A) Staining for senescence-associated β-galactosidase (SA-β-Gal) and (B) senescence-associated secretory phenotype via cytokine array (abcam, ab133997). (C) Densitometry quantification of cytokine array by Gel Analyzer on Image J and reported as fold change in average integrated pixel density compared to positive loading controls. (D) Quantitative PCR of cDNA for relative mRNA IL-6 expression levels in TLR7+ or TLR9+ MDA-MB-361 cells.

## Discussion

Consistent with observations in epithelial cells from cervical and head and neck cancers [21,22], we observed a reduction in TLR9 expression in epithelial cells from primary breast tumoral tissue compared to normal epithelium from healthy donors. Additionally, the adjacent epithelial tissue from patient matched tumoral samples trend towards a more negative immunoreactive score (IRS) for TLR9, which may suggest that BC tumors overexpress TLR9 compared to their surrounding tissue. Furthermore, these data agree with previous observations where BC epithelial cells are more TLR9+ than their adjacent tissue or stromal cells [26,33]. Yet, the opposite is true for the phenotype observed in TMA tumors, where no TLR9 positive signal was detected amongst 47 sections, despite the peritumoral tissue presenting a mean TLR9 IRS score identical to that observed in conventional tumors. One interpretation could be that conventionally sectioned tumors are larger in surface area compared to TMAs, and therefore offer a more heterogenous sampling of the tumor microenvironment. Given the BCs are prone to histological heterogeneity [6], it is possible that the full sections capture epithelial showing spatial and temporal heterogeneity [5]. In fact, the uniformity of phenotype amongst the TMA TLR9 expression supports the idea that TMA sections are a better representation of the true phenotype amongst a homogenous BC epithelial population. With regards to the timing of TLR9 expression in BC, previous work [34] in clinical samples from early noninvasive breast lesions and patients with triple negative severe oncogenesis indicate that higher TLR9 expression appears early during breast carcinogenesis, and is associated with poor long term prognosis. We posit that the loss of TLR9 expression may be a late-stage event during BC carcinogenesis.

Although high and low TLR9 expression is found in all the clinically relevant subgroupings of breast cancer, including estrogen (ER) and progesterone (PgR) receptor-positive, human epidermal growth factor receptor-2 (HER2)-positive, and triple-negative tumors (which lack the expression of all these receptors), the concurrent effect on metastasis or poor prognosis stratifies along these subcategories. For example, in TNCs, TLR9 was associated with higher expression correlating with a more fibrous and inflamed TME [30,34], and loss of this expression belies an aggressive metastasis with poor survival outlook [31,35]. In contrast, there is an inverse correlation between tumor TLR9 and ER expression [12] where ER-positive breast cancers have significantly lower levels of TLR9 expression, as compared with TNBCs and correlated with poor prognosis [25,27,34]. We observe that in our clinical samples, TLR9 is under expressed. These observations are supported by a significant reduction in TLR9 mRNA expression by microarray analysis of 59 breast cancer cell lines. Consistent with the clinical data, in our own observations of TLR9 mRNA expression on six epithelial BC cell lines, relative TLR9 expression seems to stratify according to the BC clinical subgroupings. The three TNC cell lines we analyzed (MDA-MB-231, HS578T, and Hcc1937) had the highest relative TLR9 expression compared to the Her2+ (SK-BR-3) or the ER+ (T-47D, MDA-MB-361) cell lines [36,37]. These observations may indicate a shared function of TLR9 as a biomarker with other well-known prognostic factors in BC. Furthermore, TLR9 overexpression in TLR9 positive BC cell lines increased invasiveness *in vitro* and signs of poorly differentiated tumors, particularly under stimulation with ligand CpG-DNA [13,25,26]. This may indicate that metastatic effects of TLR9 may occur in a dose dependent during BC, however the extent to which TLR9 acts as in a pro-transformative or tumor suppressant were unclear from these studies.

In head and neck cancer where endogenous TLR9 is under expressed, TLR9 exogenous expression induced cell stress (stasis in S phase), with a reported accumulation in G1 after TLR9 reintroduction [21]. Our data reflects a similar phenotype, suggesting overexpression of TLR9 may induce G1 arrest, however a more detailed analysis into whether cells are truly arrested or undergoing apoptosis is required. Furthermore, our data reflects a reduction in cell proliferation by colony formation in TLR9+ cells, which suggesting TLR9 has a specific effect on suppressing cell proliferation in BC. Furthermore, upon strong overexpression selection (i.e. increasing doses of doxycycline). To investigate if the reduced cellular proliferation was induced by cell cycle arrest, we investigated markers of cellular senescence, which can be triggered by cellular stress from the changes associated with oncogenic activation [38]. We observed a saturation of IL-6 in TLR9+ as well as IL-6-like cytokine, oncostatin-M, with a strong increase in the mRNA IL-6 profile of TLR9+ cells. IL-6 expression is reduced in invasive breast carcinoma relative to normal mammary tissue and appears to be inversely associated with histological tumor grade [39–41]. We also noted an increased secretion of GM-CSF in TLR9+ cells. Interestingly, GM-CSF is expressed in more than half of patient tumors [42] and promotes an immunosuppressive TME in BC to activate plasmacytoid dendritic cells in a regulatory Th2 [42–44]. Conversely by cytokine array, we observe a slight increase in secretion profile of TLR9+ compared to TLR7+ BC cells in VEGF, which promotes BC invasion through autocrine signaling of CXCR4 [45]. This may suggest that although TLR9+ cells are senescent, they may retain some metastatic potential. Additionally, in our study TLR9 overexpression had strong impact on secretion of CXCL-1, which is implicated in BC metastasis via recruitment of myeloid cells [46]. Collectively, these observations indicate that not only does TLR9 apparently promote senescence but seems to exhibit an immunoprofile consistent with cytokines associated with tumor suppressive microenvironments, alongside apparent changes in factors promoting invasion. Although highly indicative of a senescent mechanism, further experiments to articulate the nature of when TLR9 induces BC cell arrest and if these cells express hallmark tumor suppressor proteins like p53, p21^CIP1^ and p16^INK4A^ [47] could be helpful in elucidating the exact mechanisms of TLR9 tumor suppression in BCs. Nevertheless, between the clinical observations and those in BC cell lines, we postulate that perhaps a few key TLR9 expressing cells may induce a pro-senescent microenvironment through paracrine signaling that acts as a tumor suppressant to protect against metastasis. Furthermore, our *in vitro* data would suggest that TLR9 can act in a cell-autonomous manner in tumor suppressor function and directly impede oncogenic transformation in the epithelium of BCs.

## Materials and Methods

### Patients and samples

TLR9 Immunohistochemistry (IHC) was performed on formalin-fixed, paraffin-embedded breast tissue from King’s College Tissue Bank, UK. We randomly selected a sample size of 98 breast tissue, consisting of 29 cases of normal breast tissue, 39 cases of peritumoral breast cancer tissue, and 29 cases of invasive breast cancer tissue with their adjacent tissue. Furthermore, a tissue microarray with 49 breast cancer cases was available to us. All samples were collected at diagnosis prior to chemotherapy in compliance with and approved by the Institutional Review Boards.

### Immunofluorescence and immunohistochemistry of patient sections

Serial tissue sections were cut, deparaffinized, rehydrated, and subjected to the appropriate antigen retrieval method. The sections were then incubated with anti-TLR9 (clone D9M9H, Cell Signalling Technologies Ltd) and visualized with 3,3-diaminobenzidine and hematoxylin or with secondary anti-Rabbit-Cy3 antibody (Jackson Immunoresearch Lab). The positive tissue control consisted of archived, formalin-fixed human tonsillar and cervical tissue. Blocking peptide (produced by Cell Signalling Technologies Ltd) was used as a negative control. The samples were then examined using a light microscope. Two pathologists, blinded to the associated information about the samples, performed the scoring. The staining percentage was scored on a scale from 0-4 (0=no staining, 1=<20%, 2=20-30%, 3=40-50%, 4=50-70%, 5=>70%). The staining intensity was scored on a scale from 0-4 (0=no staining, 1=very weak, 2=weak, 3=moderate, 4=strong). We multiplied the percentage by the intensity scores to generate the immunoreactive score (IRS). The IRS score ranged from 0-15. Any IRS score above 0 was considered positive.

### Lentiviral expression vector

The human pLVUT is a lentiviral vector expressing a *Tet*-on expression cassette in a doxycycline-inducible manner. Human TLR9 and TLR7 cDNA sequence were cloned into vector as previously described [48].

### Cell Culture

MDA-MB-361 human breast cancer cells were obtained and cultured according to American Type Culture Collection (ATCC) standard protocol. Cells were maintained in Dulbecco’s modified Eagle’s medium (DMEM) supplemented with 10% fetal bovine serum. Cells were regularly tested for mycoplasma using specific primers and subcultured sterily in L2 facilities.

### Microarray analysis

TLR9 gene expression analysis was generated by using Array Studio software (Omicsoft Corporation). CEL files of BC (N=59) and immortalized (N=8) cell lines were downloaded from the Cancer Cell Line Encyclopedia CCLE [49]. Raw microarray data were processed using quantile normalization and robust multiarray average algorithm. A custom CDF file from the Brain Array database was used for the summarization (http://brainarray.mbni.med.umich.edu/Brainarray/Database/).

### RNA extraction, reverse transcriptase-PCR and qPCR

RNA was extracted using Nucleospin RNA/protein kit following the manufacturer protocol (Macherey-Nagel, Germany). Reverse transcriptase reaction was performed using 500–1000ng of RNA. For qPCR, complementary DNA were diluted 1/20 for quantitative PCR (qPCR) reactions using Mesa green qPCR Master Mix (Eurogentec, Angers, France). PCR was conducted using the Mx 3000P real-time PCR system (Stratagene, La Jolla, CA, USA). Two sets of PCR assays were conducted for each sample, the TLR9 and β2-microglobulin primers have been described [50]. Amplification specificity was assessed for each sample by melting curve analysis, and the size of the amplicon checked by electrophoresis (data not shown). Relative quantification was performed using standard curve analysis. TLR9 mRNA levels were normalized to β2-microglobulin mRNA levels and are presented as a ratio of gene copy number per 100 copies of β2-microglobulin in arbitrary units.

### IL-6 mRNA detection

IL-6 measurement was performed as described by [51].

### Immunoblot analysis

In brief, harvested cells were lysed in mild lysis buffer containing 50□mM Tris-HCl (pH 8.0), 150□mM NaCl, 1% Triton X-100, 1□mM DTT, 0.5□mM and complete protease inhibitor (Roche, Meylan, France). Cellular protein content was determined by the Bradford assay (Bio-Rad, Marnes-la-Coquette, France); used for sodium dodecyl sulphate-polyacryl amide gel and immunoblotting onto a polyvinyl difluoride membrane. After incubation with primary antibodies, proteins were detected with secondary peroxidase-conjugated antibodies (Promega, Madison, WI, USA) and ECL. All the primary antibodies for western blotting were from Cell Signaling but the β actin (MP biomedicals, Santa Ana, CA, USA).

### Proliferation curve and colony formation assay

Doubling population assay and clonogenicity assay were previously described [52], each cell type, 1×10^2^ cells were seeded into 100□μl of media in 96X microplates (E-Plate). The attachment, spreading and proliferation of the cells were monitored every 7 days for 4 weeks and counted using trypan blue dead cell exclusion. For the colony formation, cells were split for selection with doxycycline after pLVUT-TLR7/9 infection and grown as previously described [53]. Colonies were washed with PBS, fixed, and stained on the plate for crystal violet in 20% methanol.

### Cell Cycle Analysis and Senescence assays

MDAM-MB-361 overexpressing pLVUT-TLR9 or pLVUT-TLR7 conditioned in doxycycline for 3-4 days then spun at 1000g for 5 min to remove debris. The supernatant was harvested for Human Cytokine Antibody Array (abcam, ab133997) according to the manufacturer’s instructions. Dot blots were analysed as previously described [18]. Arrays were quantified using the Gel Analyzer tool on ImageJ. The average signal (integrated pixel density) was standardized to the positive loading controls for each blot. For cell cycle analysis by flow cytometry, cells were stained by Click-iT Edu (Invitrogen C10337) according to manufacturer’s instructions and analyzed on the BD Fortessa 5L. SA-β-Gal staining was performed according to the manufacture’s instructions (Cell Signaling, SA-β-Gal staining kit (#9860).

### Statistical analysis and graphics

All analyses were generated using GraphPad Prism (version 8 or later) by one-way ANOVA unless otherwise stated.

## Supporting information

original western

## Supplemental

Dox-inducible TLR9 rescue with pLVUT in MDA-MB-361. BC cell line MDAMB 361 was transduced with a doxycycline-inducible *TeT*-on expression plasmid pLVUT carrying transgene for TLR9 or TLR7 control. Cell lysates were analyzed via Western blotting 3 days post treatment. The schematic for the lentiviral vector is adapted from [54] (A). Western blot results for TLR9 and TLR7 (B).

## Statement conflict of interest

---

**No Conflict of Interest Statement**

As corresponding author, I hereby declare, on behalf of all authors of this manuscript that there are no conflicts of interest to disclose. None of the authors have received direct or indirect financial support, grants, or have any other personal relationships, interests, or affiliations that could influence or appear to influence the work reported in this paper.

Furthermore, all authors have contributed significantly to the research and preparation of this manuscript and share responsibility for its content. Any funding sources for this study have been acknowledged in the manuscript, and we confirm that the funders had no role in the study design, data collection and analysis, decision to publish, or preparation of the manuscript.

We affirm that we have no commercial or associative interest that represents a conflict of interest in connection with the work submitted.

Dr Uzma Hasan

## Notes

### Competing Interest Statement

The authors have declared no competing interest.

### Summary of Updates

removed one author as he was not involved with the work

