## Supplementary figures and images for "The Impact of TLR9 Expression Loss on Breast Cancer Tumor Cells"

### original western

TLR9 130kDa

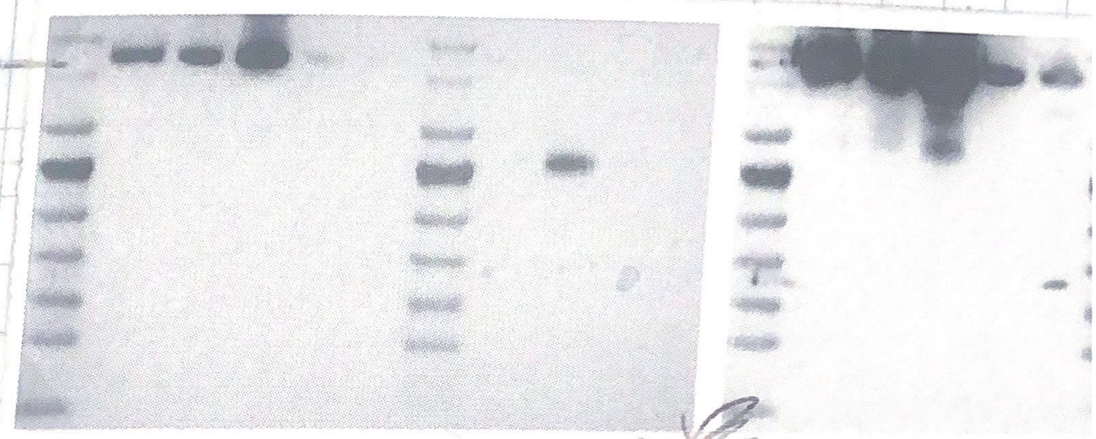

βactine

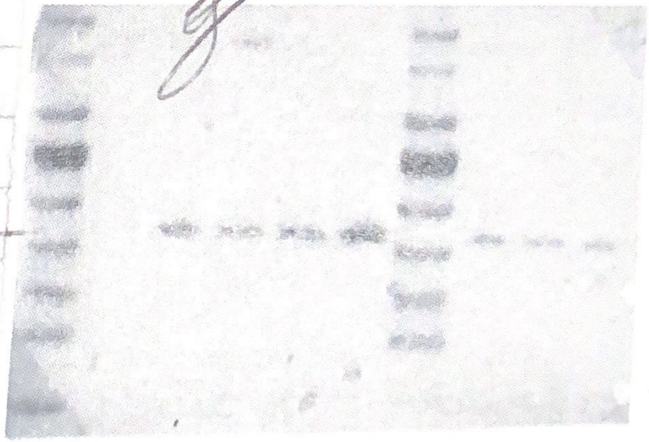

### sup 1

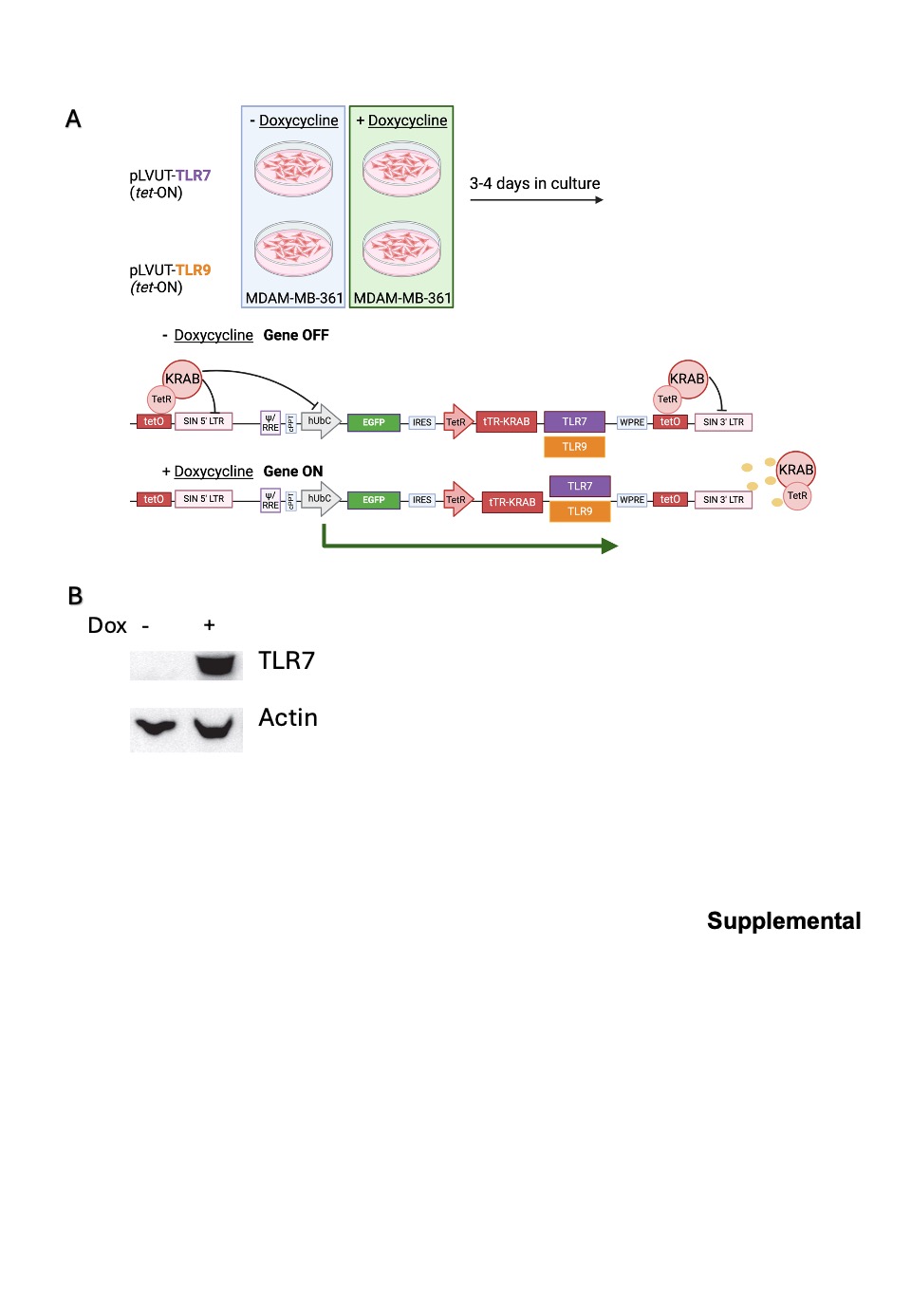
